# Parallel algorithms for phylogenetic inference under a structured coalescent approximation

**DOI:** 10.1101/2025.09.22.677844

**Authors:** Yucai Shao, Marc A. Suchard, Andrew Rambaut, Xiang Ji, Philippe Lemey, Tetyana I. Vasylyeva, Guy Baele

**Author notes:** **Corresponding author:** Guy Baele, Department of Microbiology, Immunology and Transplantation, KU Leuven, Rega Institute, Herestraat 49, 3000 Leuven, Belgium.

## Abstract

While advances in molecular epidemiology and computational modeling have enhanced our capacity to track pathogen evolution, the accurate reconstruction of spatiotemporal transmission dynamics remains essential for developing epidemic preparedness frameworks and implementing outbreak response measures. Structured coalescent models offer a phylogeographic framework by restricting lineage coalescence events to geographically proximate host populations. Although the Bayesian structured coalescent approximation (BASTA) provides a tractable approach, contemporary phylogeographic analyses involving dozens of geographic localities and hundreds to thousands of viral genomes substantially exceed the computational capacity of existing implementations. The BASTA likelihood scales cubically with deme count and quadratically with sequence count due to matrix exponentiation and pairwise coalescent probability calculations. Here, we introduce a comprehensive algorithmic restructuring of the structured coalescent likelihood that eliminates redundancies, optimizes memory access, and exposes parallelization opportunities. Our approach reorganizes computations along three dimensions: (i) independent calculation of deme-transition probability matrices across time intervals; (ii) simultaneous evaluation of partial likelihood vectors within temporal slices; and (iii) concurrent aggregation of coalescent probabilities. Algorithmic restructuring cuts average coalescent likelihood computation by 7–8 fold, and parallelization further boosts performance to 10–26 fold, enabling joint phylogeographic analyses of dengue virus across 10 South American countries and H5N1 avian influenza across 20 Eurasian regions to finish in a fraction of prior time. This computational efficiency also enables comparison between backward-in-time structured coalescent approximations and forward-in-time phylogeographic methods, revealing that the former provides appropriately conservative posterior estimates, particularly at intermediate phylogenetic depths. We integrate our implementation into the popular BEAST X and BEAGLE software packages, with an accompanying interface in BEAUti X to easily set up the analyses, providing researchers with an accessible and scalable tool for real-time phylogeographic surveillance of rapidly evolving pathogens.

## 1 Introduction

Structured coalescent models provide a sophisticated mathematical framework for understanding genealogical processes in spatially subdivided populations. These models extend classical coalescent theory by explicitly incorporating migration between discrete subpopulations (demes), enabling researchers to investigate how population structure influences genetic diversity and to reconstruct complex demographic histories from sequence data [Nordborg, 1997]. By integrating coalescent theory with explicit spatial structure, structured coalescent models enable rigorous statistical inference of demographic parameters and migration histories. These approaches are particularly crucial when population subdivision substantially influences genealogical patterns [Donnelly and Tavaré, 1995, Notohara, 1990, Wakeley and Aliacar, 2001].

Lemey et al. [2009] demonstrated the necessity of spatial structure for accurate phylogeographic reconstruction of H5N1 avian influenza, revealing complex interregional transmission dynamics that would be obscured under panmictic assumptions. Bedford et al. [2010] quantified how geographic compartmentalization shapes influenza A/H3N2 evolution, with migration rates between regions creating measurable constraints on viral genetic diversity. Beyond pathogens, structured coalescent frameworks have demonstrated how glacial isolation in refugia shaped genetic structure and migration patterns in alpine plants and other taxa [Eme et al., 2023]. These studies highlight scenarios where failing to account for population structure leads to substantial misinterpretation of evolutionary processes and demographic history.

The exact structured coalescent model [Hudson et al., 1990] provides a comprehensive framework for studying genealogies across subdivided populations by explicitly accounting for both the tree structure and individual migration events. While calculating the likelihood is straightforward when migration events are fully known, in practice these events are typically unobserved. For example, in epidemiological studies, we might observe viral sequences from individuals with known travel histories to endemic regions, but the exact timing and number of introduction events that gave rise to local transmission chains remain unknown. This uncertainty necessitates integration over all possible migration histories, which can be addressed by sampling the tree annotated with migration events using structured tree operators [Vaughan et al., 2014]. However, this requirement renders the exact model computationally intractable for large datasets. To address this computational challenge, structured coalescent approximation (SCA) models have been developed [Palczewski and Beerli, 2013, Müller et al., 2017, 2018], with the Bayesian structured coalescent approximation (BASTA) [De Maio et al., 2015b] representing a prominent approach. The exact structured coalescent model accounts for correlations among lineages that coexist in the phylogeny by explicitly modeling the joint probability of the phylogenetic tree and the configurations of all lineages across possible states as they change over time. BASTA introduces computational tractability by assuming independence between lineage states and the coalescent process intervening between events. This approximation enables the calculation of partial likelihood vectors through a backward-in-time (BIT) continuous-time Markov chain (CTMC) process, characterized by a transition rate matrix solely determined by migration rates. BASTA employs a trapezoidal rule-based approximation for likelihood calculations, streamlining the integration process over intervals.

In contrast, forward-in-time (FIT) CTMC models for discrete trait phylogeography [Lemey et al., 2009] simplify the modeling process by assuming complete independence between migration events and the tree generation process. This leads to a joint likelihood that is computationally simpler but potentially less accurate in scenarios when sampling is uneven or biased toward specific populations [De Maio et al., 2015b].

Phylogeographic reconstruction under structured coalescent models is expected to perform within reasonable computational limits when few subpopulations are present [De Maio et al., 2015b]. However, the ability to accommodate larger numbers of demes is increasingly important as local-scale analyses require fine-grained resolution between smaller administrative units such as cities or districts to reveal transmission patterns relevant for targeted public health interventions. The computational burden of existing structured coalescent implementations scales poorly with the number of demes, becoming prohibitive for practical inference as this number increases. This limitation not only restricts our ability to fully exploit rich, multi-scale datasets but also raises parameter identifiability concerns when many demes are considered simultaneously. Consequently, practical analyses using BASTA typically require the assumption that all deme sizes are equal and constant over time to avoid over-parameterization. Even with this restriction in place, Layan et al. [2023] found that BASTA proved impractical to perform discrete phylogeography on datasets consisting of 500 sequences that were sampled from seven demes, both as a result of extremely high calculation times (over 70 hours per million iterations in the original implementation) and of convergence issues due to insufficiently high effective sample size (ESS) values after a long run time and multi-modality in the model’s continuous parameters.

Earlier efforts to harness parallel computing in phylogenetics have demonstrated significant speedups using both multi- and many-core architectures [Suchard and Rambaut, 2009, Baele et al., 2017, Gangavarapu et al., 2024]. Further, the continued development of powerful hardware architectures has resulted in markedly faster phylogenetic and phylogeographic inference through the use of high-performance computing libraries. However, structured coalescent models have remained computationally intractable for large datasets despite these advances.

In this work, we overcome this limitation through a dual strategy: first, we fundamentally reformulate BASTA’s likelihood computation within BEAST X [Baele et al., 2025] to eliminate redundant calculations and optimize memory access patterns, achieving 7-8-fold speedups even in serial execution (detailed benchmark results can be found in Supplementary Materials). Second, we leverage this reformulated algorithm to enable multi-threaded execution across available central processing unit (CPU) cores by extending the high-performance BEAGLE library [Suchard and Rambaut, 2009, Ayres et al., 2019]. We partition the structured coalescent likelihood workload to enable concurrent processing, distributing indepen-dent tasks across threads and minimizing synchronization overhead. The algorithmic reformulation alone dramatically improves computational efficiency, while the subsequent parallelization yields aggregate speedups of 10-to 26-fold. As a result, our framework renders previously intractable structured coalescent analyses computationally feasible, enabling researchers to analyze substantially larger datasets and perform more complex tasks.

## 2 Methods

In the SCA framework, the structured coalescent density calculations are broken down by intervals, each defined by coalescent or sampling events within the phylogenetic tree. Within each coalescent time interval, the structured coalescent likelihood contributions for internal nodes can be computed independently. For each node in the interval, these computations involve updating both the partial likelihood vectors based on the node’s descendant states and the transition probabilities over the interval. Because these updates do not depend on the simultaneous processing of other nodes within the same interval, we can parallelize the computations across processor cores. This design enables efficient multi-threaded evaluation of the structured coalescent likelihood, as illustrated in Figure 1 and detailed in the following section. We here describe a reformulation of the structured coalescent density computation to enable parallel processing on multi-core CPUs, by organizing the computations into independent node-specific operations that can be carried out concurrently.

**Figure 1.**
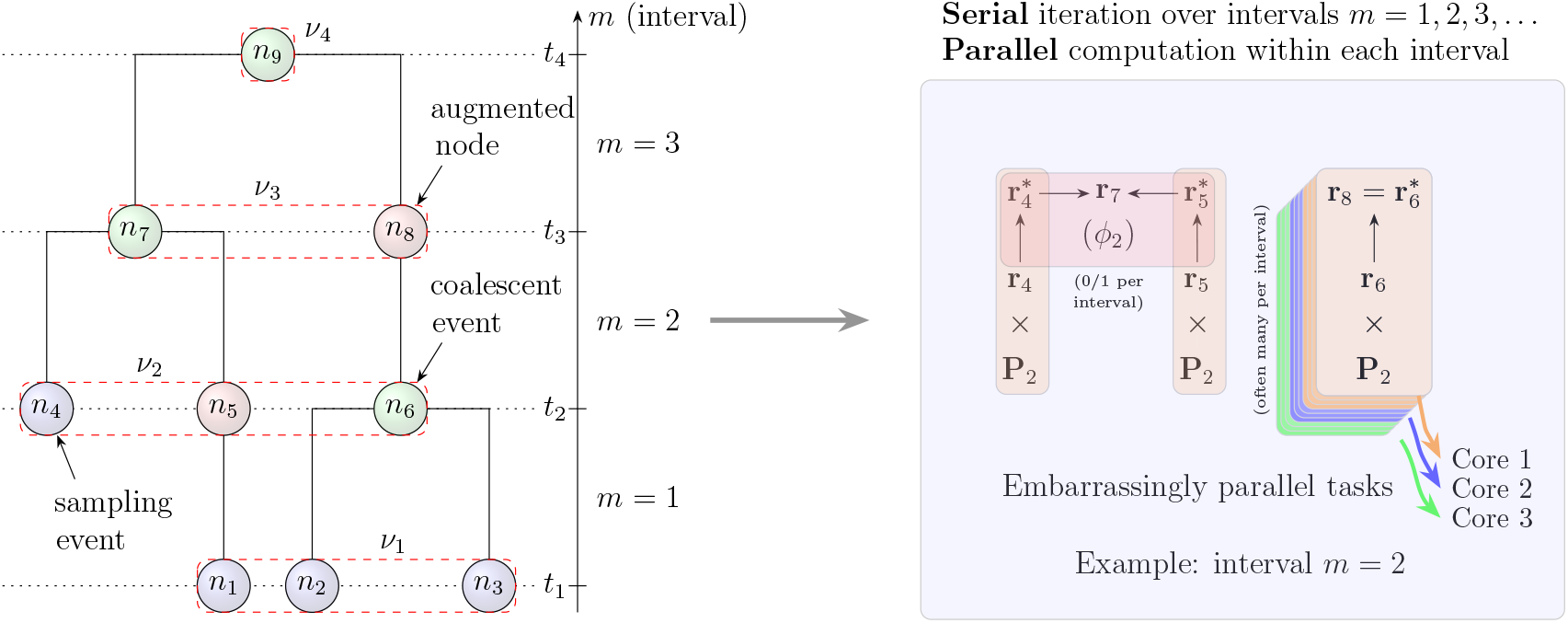
Illustration of augmented nodes and coalescent intervals in structured coalescent likelihood computations. The left panel shows an augmented phylogeny with degree-2 nodes inserted at time boundaries, creating intervals *m* = 1 through *m* = 3 between times *t*_1_ and *t*_4_, with corresponding node sets *ν*_1_ through *ν*_4_. The right panel depicts the parallel computation scheme for interval *m* = 2, where nodes within an interval are processed independently across multiple CPU cores, while intervals are processed sequentially.

Let ℱ denote a rooted, bifurcating phylogenetic tree representing the evolutionary relationships among *N* sampled sequences from *S* demes. The tree consists of tip nodes corresponding to sampling events and internal nodes representing coalescent events where lineages merge backward in time (see Figure 1). Each node *i* ∈ ℱ is associated with a time point and a location state within a structured population of demes. We define a sequence of time points **T** = (*t*_1_, *t*_2_, …, *t*_*M*+1_) where *t*_1_ *< t*_2_ *<* … *< t*_*M*+1_, ordered from present to past. Here, *t*_1_ represents the most recent sampling time and *t*_*M*+1_ represents the time of the root.

The integer *M* denotes the number of time intervals in the tree, where *M* ≤ 2*N* − 1. Each time point *t*_*i*_ corresponds to either a sampling event (for tip nodes) or a coalescent event (for internal nodes). To facilitate computation, we augment ℱ by inserting degree-2 nodes at all points where branches intersect time slice boundaries between consecutive time points in **T**. These augmented nodes have exactly one parent and one child (e.g., node *n*_8_ in Figure 1). This augmentation partitions the tree into *M* discrete time intervals: *m*_*i*_ = [*t*_*i*_, *t*_*i*+1_) for *i* = 1, 2, …, *M* .

Within each interval *m*_*i*_, we compute partial likelihood vectors **r**_*i*_ for all nodes *i* (both original and augmented) that exist within that interval. These vectors are propagated recursively through the tree structure to enable efficient phylogenetic likelihood calculations. Similar to Felsenstein’s pruning algorithm [Felsenstein, 1981], **r**_*i*_ = (*r*_*i*1_, …, *r*_*iS*_) quantifies the probability of all location data ⌊**X**_*i*_⌋ below (or extant to) node *i* in ℱ given that the CTMC process realizes state *s* at node *i*. In our notation, *S* denotes the total number of discrete geographic states (i.e., demes) over which taxa are distributed. However, unlike the standard pruning algorithm, our approach accounts for coalescent events (such as at nodes *n*_6_ and *n*_7_) that occur between lineages in different demes with (inverse) population sizes ***λ*** (see Figure 1).

The SCA recursion differs from traditional pruning in three key ways. First, it traverses nodes in *reverse-level-order*, processing all nodes within an interval (e.g., all nodes in *ν*_1_ in Figure 1) before moving to earlier intervals. Second, it computes additional quantities for each interval to trace the coalescent process underlying ℱ. Third, this interval-based decomposition exposes significant parallelization opportunities both within node-specific calculations and across all nodes in an interval. Starting from ℱ, we begin our computation by iterating over inter-coalescent intervals with indices *m* = 1, …, *M* . Within each of these dependent iterates, we first collect the set of nodes that lie at *t*_*m*_ into *ν*_*m*_ and define the inter-coalescent interval length *τ*_*m*_ = *t*_*m*+1_ − *t*_*m*_. For intervals where *m* ≤ *M*, we compute the finite-time transition matrix **P**_*m*_ = exp (*τ*_*m*_**Q**), where **Q** is the infinitesimal rate matrix (generator) of the CTMC governing the migration process between demes. This matrix **P**_*m*_ characterizes the state-change probabilities for lineages transitioning between demes during the time interval *τ*_*m*_. For each node *i* ∈ *ν*_*m*_ (potentially in parallel), we propagate

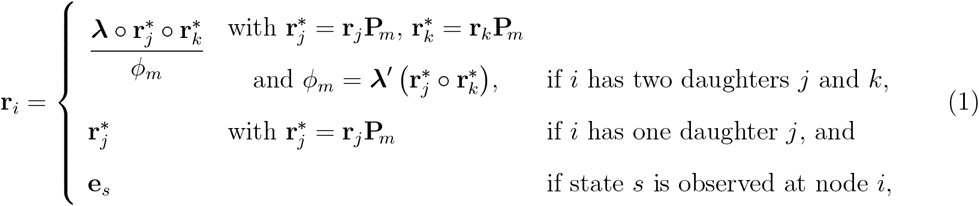

where ∘ is the Hadamard product and **e**_*s*_ is the standard unit vector in the direction of *s*. If no coalescent event occurs in the interval, we set *ϕ*_*m*_ = 1. Since at most one coalescent event can occur, at most two propagated partial likelihood vectors 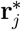 and 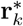 for the event represent unique values and benefit from caching along with *ϕ*_*m*_.

The right panel of Figure 1 illustrates how this augmentation and interval-based decomposition creates natural parallelization opportunities. Within each interval, nodes can be processed independently as demonstrated for interval *m* = 2. Here, partial likelihood vectors for nodes *n*_4_, *n*_5_, and *n*_6_ are propagated through matrix-vector multiplications 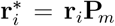 as separate operations assigned to different CPU cores (shown in green, blue, and orange), while the coalescent event at node *n*_7_ combines propagated vectors 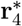 and 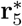 to compute both the new partial likelihood and coalescent contribution *ϕ*_2_. This interval-based parallelization strategy, where all nodes within an interval are processed in parallel before moving to the next interval, efficiently utilizes modern multi-core architectures.

Once completing the traversal, the SCA log-likelihood becomes

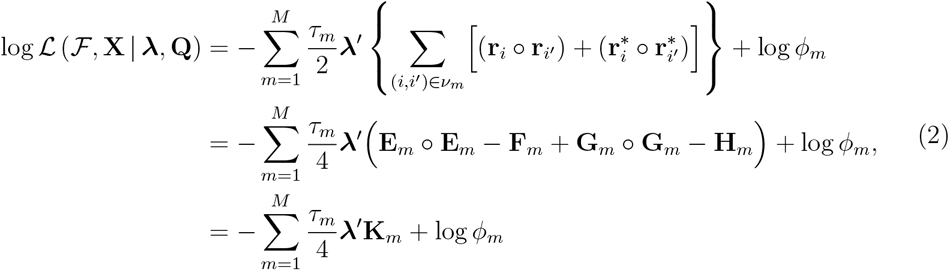

where (*i, i*^*′*^) ∈ *ν*_*m*_ iterates over all unique node pairs, and *S* × 1 transformation vector **K**_*m*_ = **E**_*m*_ ∘ **E**_*m*_ − **F**_*m*_ + **G**_*m*_ ∘ **G**_*m*_ − **H**_*m*_ and intermediate vectors:

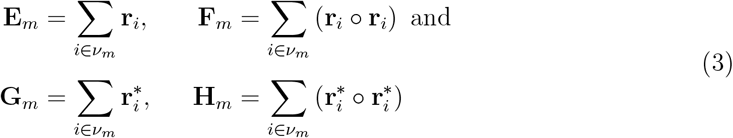

stream-line computational work into simple (parallelizable) reductions over all nodes in a coalescent interval. Specifically, it is feasible, and indeed our practice, to pre-compute all transition matrices, **P**_*m*_, for each interval *m* concurrently. The probability vectors **r**_*i*_ and 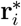 computed during the tree traversal are stored for subsequent reuse in calculating these intermediate summation terms in Equation 3 and performing ancestral state reconstruction, thereby avoiding redundant recomputation across these procedures.

Our methodology iterates through three core steps: the pre-computation of **P**_*m*_, the peeling process for updating all **r**_*i*_, and the reductions within the intervals to the intermediate vectors. The computational complexity of these steps can be characterized by the following orders: the computation of the transition matrices operates at 𝒪(*NS*^3^), the evaluation of the partial likelihood vectors for all nodes extant at *M* interval time points is 𝒪(*N* ^2^*S*^2^), while the reduction within each interval is 𝒪 (*N* ^2^*S*).

The computational complexities identified above present significant challenges for analyzing contemporary phylogeographic datasets. To transform these theoretical complexity bounds into practical computation times, we developed a novel parallel computing framework that exploits the inherent independence within each of the three core steps. We implemented our CPU parallelization using OpenMP (Open Multi-Processing), a shared-memory threading API, to target the three key computational phases identified above. We now detail how parallelization strategies specifically target each computational phase to achieve substantial speedups on modern multi-core architectures.

### Transition probability matrix computation

The SCA framework requires computing finite-time transition probability matrices **P**_*m*_ = exp(*τ*_*m*_**Q**) for each inter-coalescent interval *m*. Matrix exponentiation via eigen decomposition dominates the computational cost at 𝒪(*S*^3^) per matrix [Moler and Van Loan, 2003]. We exploit the statistical independence between transition matrices to implement a parallelization strategy. Since each interval’s transition matrix depends only on its duration *τ*_*m*_ and the shared rate matrix **Q**, we distribute the *M* transition matrix computations across *P* available processor cores. This achieves speedup that scales nearly linearly with the number of processor cores when *M* ≫ *P* . The parallelization reduces the effective complexity to 𝒪 (*NS*^3^*/P* ) while maintaining numerical stability through established matrix exponentiation algorithms.

### Parallel peeling updates within intervals

The peeling algorithm propagates partial likelihood vectors backward through the augmented phylogeny via post-order traversal. Within each coalescent interval *m*, we must compute 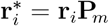 for all nodes *i* ∈ *ν*_*m*_. This matrix-vector multiplication exhibits complexity 𝒪 (*S*^2^) per node, yielding aggregate complexity 𝒪 (|*ν*_*m*_| *S*^2^) per interval.

The statistical independence of lineage state transitions within intervals enables embarrassingly parallel computation. We partition the node set *ν*_*m*_ among *P* threads using dynamic scheduling to ensure proper load balance. Each thread computes 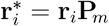 for its assigned subset of nodes without inter-thread communication or synchronization. For intervals where |*ν*_*m*_| ≫ *P*, the speedup scales nearly linearly with processor count, approaching the ideal speedup of *P*, as each processor handles |*ν*_*m*_|*/P* independent matrix-vector multiplications. However, when an interval contains few lineages relative to the number of available processors, the overhead of thread creation and synchronization may exceed the computational benefit of parallelization. In such cases, our implementation adaptively reduces the number of active threads or processes these intervals serially to maintain computational efficiency. This adaptive strategy ensures high parallel efficiency throughout the BIT traversal, from recent intervals with numerous extant lineages to ancient intervals with fewer lineages approaching the root.

### Parallel reductions across intervals

The SCA model density computation requires aggregating contributions across all coalescent intervals according to Equation 3. Each interval *m* contributes terms involving pairwise products of partial likelihood vectors and potential coalescent events. Specifically, we must compute the intermediate vectors **E**_*m*_, **F**_*m*_, **G**_*m*_, and **H**_*m*_ defined in Equation 2, followed by the transformation vector **K**_*m*_ that aggregates these contributions.

We implement a two-stage parallel reduction strategy. In the first stage, we exploit the independence of interval-specific calculations by distributing intervals among threads. Each thread computes the intermediate vectors and transformation vector for its assigned intervals, accumulating partial log-likelihood contributions locally. This parallelization transforms the 𝒪 (*N* ^2^*S*) sequential computation into 𝒪 (*N* ^2^*S/P* ) parallel work, achieving speedup proportional to the number of processors when the number of intervals exceeds *P* . The second stage aggregates partial results from all threads into the global log-likelihood. We implement this reduction using OpenMP atomic operations to ensure numerical correctness under concurrent updates. To minimize contention, each thread maintains a private accumulator for its assigned intervals, performing a single atomic addition to the global sum upon completion. This design confines synchronization to 𝒪 (*P* ) atomic operations, negligible compared to the 𝒪 (*N* ^2^*S*) arithmetic operations.

### Scalability and limitations

Our multi-level parallelization yields substantial theoretical throughput improvements, though practical speedups encounter limits defined by Amdahl’s Law, which states that the maximum speedup of a parallel algorithm is constrained by the fraction of the computation that remains serial [Amdahl, 1967]. Ideally, fully parallelized phases scale roughly proportionally to 1*/P* (number of CPU cores). However, a key limitation arises from the peeling process itself, which is executed via a serial loop over intervals backward in time. This serial loop introduces a non-trivial sequential component, constraining scalability as each interval must be processed fully before moving on to the next. Additionally, overheads from thread coordination and aggregation introduce further small serial components, such as combining interval results within critical sections or handling intervals with insufficient work-loads to justify parallelization. According to Amdahl’s Law, these serial fractions eventually dominate runtime as *P* increases, causing diminishing returns when adding more threads. Moreover, memory bandwidth and cache performance impose additional constraints, while simultaneous memory accesses by multiple cores to shared data structures (e.g., large state probability vectors and transition matrices) can saturate memory bandwidth, limiting scaling. Nonetheless, most calculations (e.g., matrix exponentiation and multiplications) are computationally intensive with good cache locality, making our approach predominantly compute-bound rather than memory-bound for datasets with hundreds of *N* and tens of *S*. Beyond certain core counts, scaling efficiency tapers off due to thread-management overhead and minor serial steps. By parallelizing at the thread level, we achieve near-optimal CPU resource utilization, significantly reducing iteration runtime complexity to approximately 𝒪 (*NS*^3^*/P* + *N* ^2^*S*^2^*/P* + *N* ^2^*S/P* ) plus minimal serial overhead. In practice, our multi-core CPU implementation enables SCA analyses on datasets and models that are orders of magnitude larger and more complex than existing methods can handle, effectively combining the theoretical accuracy of structured coalescent models with the computational feasibility required for real-world applications.

## 3 Results

### 3.1 Benchmark Result

We evaluate the performance gains afforded by the parallel algorithms presented in this paper using two distinct computing systems and three phylogeographic datasets of varying size and complexity. System A is a high-performance computing cluster node with dual Intel Xeon Silver processors (48 cores total) running at 2.20GHz and 512GB DDR4 RAM. System B is a local system equipped with a 24-core Intel processor running at 5.60GHz and 64GB DDR4 RAM.

Our benchmark datasets include: (i) European bat lyssavirus type 1 (EBLV), characterized by 51 nearly complete genome sequences obtained from serotine bats (Eptesicus serotinus) sampled across 3 distinct European regions (France, Spain, and the Netherlands) over a 45-year period, focusing on viral dispersal patterns associated with bat population structure [Troupin et al., 2017]; (ii) zika virus (ZIKV), featuring 283 viral genomes collected primarily from infected travelers between 2013 and 2018, capturing virus transmission dynamics across 22 countries during the epidemic peak in the Americas [Grubaugh et al., 2019]; and (iii) porcine epidemic diarrhea virus (PEDV), comprising 756 sequences collected from 26 global regions, including significant sampling within China between 2011 and 2020, where discrete phylogeographic analyses revealed associations with swine trade patterns [He et al., 2022]. This diverse collection enables us to evaluate performance across phylogenetic trees with different topologies, branch lengths, and state-space dimensionality.

We conducted performance benchmarks by measuring structured coalescent likelihood calculation times during actual Markov chain Monte Carlo (MCMC) sampling runs with fixed phylogenetic trees. To isolate the computational burden of the structured coalescent model, we fixed the tree topology, branch lengths, and population sizes while exclusively updating the migration rate matrix parameters during each MCMC iteration. This focused approach necessitates recalculation of transition probability matrices, subsequent partial likelihood vector propagation, and final likelihood aggregation, collectively constituting the primary computational steps in structured coalescent inference. All timing measurements were averaged over 400 MCMC iterations to provide robust estimates of computational performance across different implementation strategies.

On System A (Figure 2 upper panel), our parallel BEAGLE implementation achieves maximum speedups of approximately 15× for EBLV, 26× for PEDV, and 22× for ZIKV compared to the BASTA package in BEAST 2.7.7 [De Maio et al., 2015b]. Notably, optimal performance is typically reached with 8 to 16 CPU threads, with performance degradation observed when scaling to 36 or 48 threads, particularly for the ZIKV dataset where speedup factors drop below 4× and 3×, respectively.

**Figure 2.**
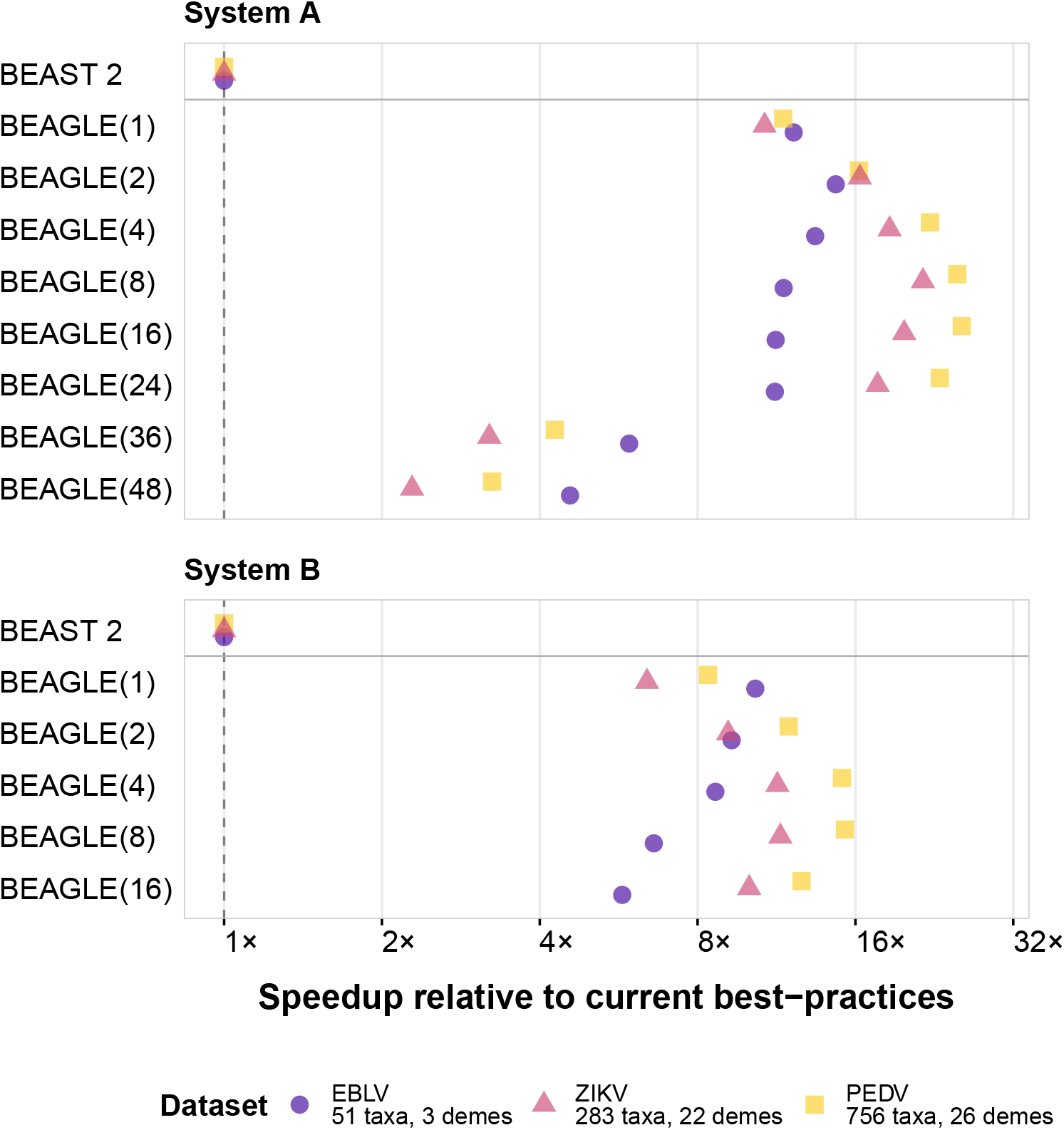
Performance comparisons between structured coalescent implementations in BEAST 2.7.7 and BEAST X for SCA model inference. Speedup factors are relative to the BASTA package in BEAST 2.7.7 for average likelihood calculation time across three datasets (EBLV, PEDV, and ZIKV) on two distinct computing systems. BEA-GLE(n) denotes the BEAGLE CPU implementation using *n* cores (number in parentheses). For our largest and most complex dataset (PEDV), BEAST X with BEAGLE(16) achieves a performance increase of up to 26× over BEAST 2.7.7 on System A, highlighting significant computational improvements for extensive phylogeographic analyses.

System B (Figure 2 lower panel) shows similar performance patterns, with maximum speedups of approximately 10× for EBLV, 15× for PEDV, and 12× for ZIKV. In this system, the optimal thread count varies by dataset. For EBLV, the smallest dataset, non-threading implementation offers nearly the maximum attainable speedup (10.3×), suggesting that for small datasets, the overhead of parallel processing may offset its benefits beyond a certain point. The PEDV dataset shows the best performance with 8 CPU threads (15× speedup), while ZIKV benefits most from 4-8 threads (11-12× speedup).

An important observation across both systems is the diminishing returns or even performance degradation when using thread counts beyond the optimal number for a given dataset. This is consistent with increased thread management overhead and memory bandwidth lim-itations. The relationship between dataset complexity and optimal thread count suggests that larger state spaces (more taxa from more demes) benefit more from parallelization up to a certain point, after which resource contention becomes limiting.

### 3.2 Scaling Experiment

To quantify how our parallelization approach transforms algorithmic scaling behavior, we conducted controlled experiments across two critical dimensions of structured coalescent computation: location states (*S*) and sequences (*N* ). Starting from the full 769-taxon PEDV dataset with associated demes, we generated reduced datasets at *N* = 128, 256, and 512 by sampling at least one taxon per deme and weighting additional samples according to the observed state frequencies. Likewise, to vary model complexity, we regrouped the sampled taxa into *S* = 2, 4, 8 and 16 demes by relabeling taxon state attributes.

Figure 3 shows semi-logarithmic plots depicting runtime scaling for both single-threaded and parallel BEAGLE implementations with 8 cores on the PEDV dataset. In this visualization, straight lines indicate exponential scaling, pronounced curvature reflects higher-order polynomial growth, and flatter trajectories suggest improved efficiency with lower complexity. The comparison clearly demonstrates that our parallel implementation significantly flattens these curves across increasing *N* and *S*, illustrating a meaningful transformation in computational complexity.

**Figure 3.**
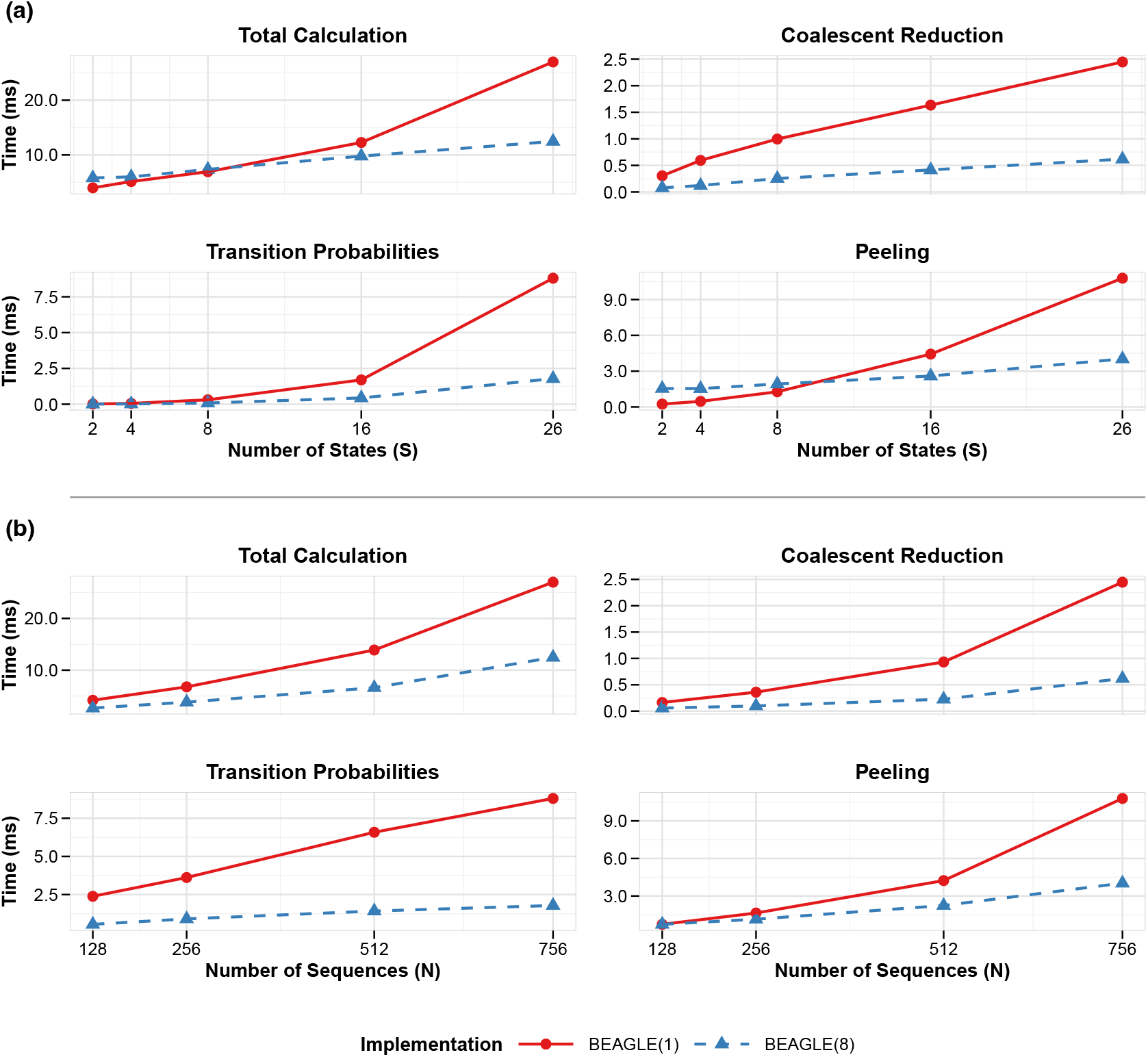
Computational scaling of structured coalescent approximation operations in BEAGLE with varying parallelization. (a) Scaling with respect to the number of states (*S*), showing performance of four key operations: Total Calculation (the overall SCA likelihood computation), Coalescent Reduction (the marginalization over coalescent configurations), Transition probabilities (computation of CTMC transition matrices between states), and Peeling (the post-order traversal for computing partial likelihoods along branches); (b) scaling with respect to the number of sequences (*N* ), demonstrating how the same operations scale with increasing phylogenetic tree size; Both panels compare single-threaded BEAGLE(1) versus 8-threaded BEAGLE(8) implementations. All timing measurements represent the average time per operation over 200 iterations.

For location state scaling (Figure 3a), the single-threaded implementation shows increasingly steep trajectories as *S* grows, particularly in the transition probability calculations where the curve’s pronounced upward curvature reflects its theoretical O(*S*^3^) complexity dominance. In contrast, our 8-thread implementation demonstrates a markedly gentler curve, indicating that our multi-level parallelization successfully mitigates the impact of the highest-order complexity terms. This effect becomes most apparent beyond *S* = 8, where the trajectories further diverge. Similarly, for sequence scaling (Figure 3b), the operations display characteristic polynomial growth patterns with varying curvature degrees. The peeling operation demonstrates the most dramatic complexity reduction, with the single-threaded implementation showing a steep trajectory consistent with its theoretical 𝒪(*N* ^2^*S*^2^) complexity, while the parallel implementation maintains a substantially flatter curve approaching the optimized 𝒪(*N* ^2^*S*^2^*/P* ) scaling. This change in the curve trends – and not merely vertical displacement – confirms that our parallelization strategy achieves more than constant-factor speedups; it effectively transforms the algorithm’s practical complexity class by reducing the growth rate coefficients of the dominant terms.

## 4 Real World Examples

To demonstrate the practical utility of our parallel algorithms for the SCA model, we analyzed two viral datasets that have been previously studied using discrete phylogeographic methods. The computational speedup achieved by our implementation makes SCA model inference considerably more practical for these datasets, which were previously computationally intensive and time-consuming to analyze. We adopt a Metropolis-within-Gibbs approach [Tierney, 1994] and use a random-scan Gibbs cycle to systematically cycle through the sampling of the phylogenetic tree topology, branch lengths, evolutionary rate parameters, and subsequently other model components. For all analyses, we discarded 10% of the MCMC samples as burn-in.

### 4.1 Dengue virus serotype 1 in Brazil

We analyzed the spatial dynamics of dengue virus serotype 1 (DENV-1) across Brazil and neighboring South American countries using the dataset from Nunes et al. [2014]. This dataset comprises 287 full genome sequences from 10 locations, including 42 Brazilian sequences that form three distinct monophyletic lineages within genotype V. The original study identified multiple introductions of DENV-1 into Brazil and demonstrated that air travel and/or mosquito vectors, rather than *Aedes aegypti* infestation levels or geographic distance, drove viral spread within the country.

Following previous discrete phylogeographic methodologies [Lemey et al., 2009, Edwards et al., 2011], we applied consistent evolutionary models for both the forward-in-time CTMC [Lemey et al., 2009] and backward-in-time SCA implementations to ensure comparability. For the nucleotide substitution process, we utilized the HKY model [Hasegawa et al., 1985] with rate variation across sites modelled using a discretized gamma distribution (HKY+Γ) [Yang, 1994], placing a log-normal prior (mean = 1.0, standard deviation = 1.25 in log space) on the transition-transversion rate ratio (*κ*) and an exponential prior (mean = 0.5) on the shape parameter (*α*) of the gamma distribution. To accommodate evolutionary rate variation across lineages, we implemented an uncorrelated relaxed molecular clock with an underlying log-normal distribution [Drummond et al., 2006]. The mean evolutionary rate received a CTMC rate-reference prior [Ferreira and Suchard, 2008] to improve mixing, while we assigned an exponential prior with mean 1*/*3 to the standard deviation of the lognormal distribution. This parameterization allows for substantial rate heterogeneity while maintaining computational efficiency.

For the discrete phylogeographic component, we employed an asymmetric substitution model across the 10 geographic locations, with gamma-distributed priors (shape = 1.0, scale = 1.0) on 90 directional migration rates. Since the FIT approach assumes equal effective population sizes across all demes [Lemey et al., 2009], we fixed all population sizes to be equal to ensure compatibility between the FIT and BIT implementations, following established practice in comparative phylogeographic studies [De Maio et al., 2015a, Müller et al., 2018]. The MCMC analysis ran for 200 million iterations for the BIT model, sampling every 10 000 iterations. We assessed convergence using Tracer v1.7 [Rambaut et al., 2018], ensuring that all parameters achieved ESS values greater than 100 and that trace plots showed adequate mixing. We constructed summary trees using the highest independent posterior subtree reconstruction (HIPSTR) approach in TreeAnnotator X [Baele et al., 2024].

Our parallelized implementation for the BIT model in BEAST X achieved a remarkable speedup compared to its standard structured coalescent implementation in BEAST 2.7.7. To achieve statistically robust posterior distributions with ESS values exceeding 100 for all parameters, the original structured coalescent implementation in BEAST 2.7.7 required over seven days of continuous computation, whereas our parallel algorithms using two processor cores achieved equivalent ESS values in approximately one and a half days.

The phylogeographic reconstructions reveal a complex history of DENV-1 circulation in South America (Figure 4). Both the FIT and BIT methods identify three major Brazilian lineages, each representing an independent introduction event. The earliest lineage emerges in the mid-1980s, coinciding with the re-establishment of dengue in Brazil after decades of absence. The phylogenetic structure clearly shows this lineage persisting and diversifying within Brazil through the 1990s, with occasional spread to neighboring countries including Argentina and Paraguay. A second introduction occurs approximately a decade later in the mid-1990s, again showing primarily Brazilian circulation with limited regional spread. The third and most recent introduction appears around 2006, representing the smallest lineage in terms of sampled diversity.

**Figure 4.**
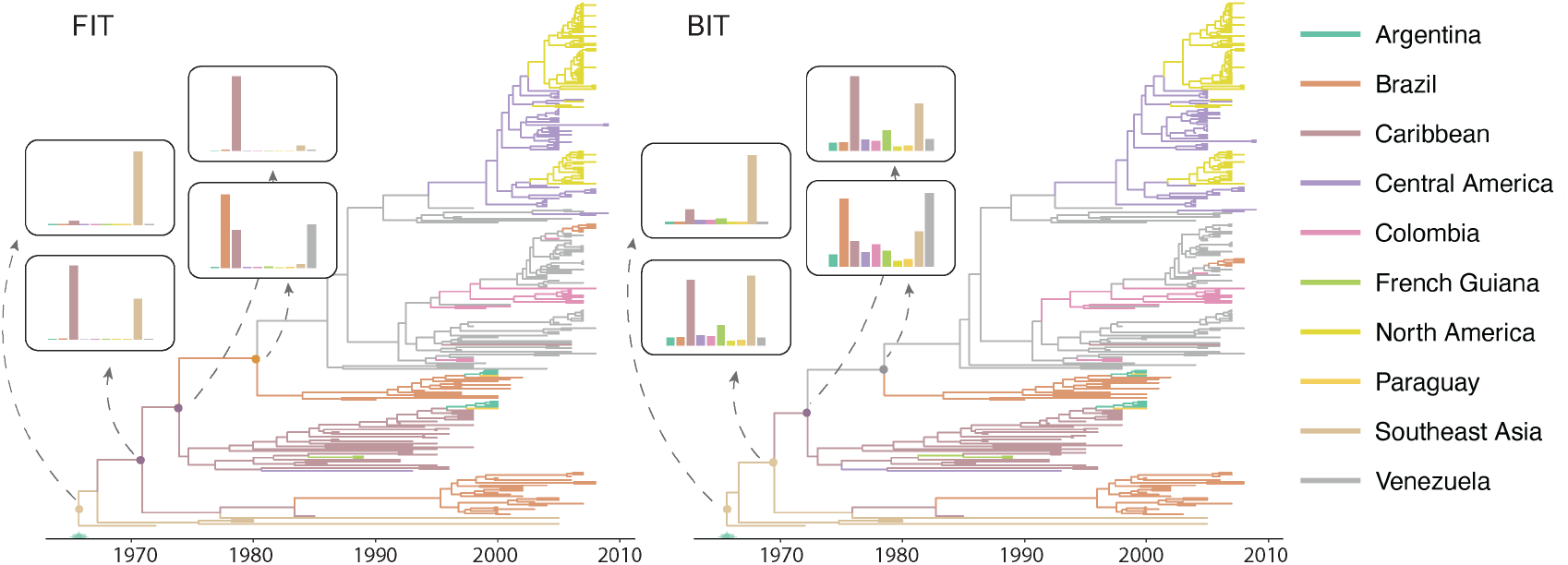
Phylogeographic reconstruction of dengue virus serotype 1 (DENV-1) showing three distinct Brazilian lineages. Highest independent posterior subtree reconstruction (HIP-STR) trees with branches colored according to the most probable location state for both FIT and BIT implementations. Insets show the posterior distribution of ancestral location states for key nodes along the Brazilian lineages. The BIT method shows broader posterior distributions across locations compared to FIT, particularly for intermediate nodes. Green bar on the time axis represents the 95% HPD credible interval for the root age.

A striking feature of the comparison between FIT and BIT methods lies in their treatment of ancestral location uncertainty. The FIT approach tends to concentrate posterior probability on single locations at internal nodes, often showing near-complete support for either Caribbean or Brazilian ancestry. In contrast, the BIT method consistently produces more diffuse posterior distributions, acknowledging substantial uncertainty about ancestral locations. This difference is particularly pronounced at intermediate nodes ancestral to Brazilian clades, where BIT assigns non-negligible probability across the Caribbean and northern South America, as well as Southeast Asia while FIT tends toward point estimates. For instance, at the basal nodes of the major Brazilian lineages, FIT assigns overwhelming support to Caribbean origins, while BIT – though still distributing a substantial portion of probability to the Caribbean – also assigns notable probability to Venezuela, other South American countries and Southeast Asia.

The broader uncertainty profiles from the BIT method may provide a more realistic assessment of what can reliably be inferred from the available data, particularly given the sparse and opportunistic nature of viral sampling across South America during this time period. The fact that both methods converge on similar conclusions regarding the timing and general source regions of introductions, despite their different uncertainty assessments, strengthens confidence in these broad epidemiological patterns.

### 4.2 Avian influenza A H5N1 hemagglutinin across Eurasia

We analyzed the spatial diffusion of highly pathogenic avian influenza A-H5N1 using the hemagglutinin (HA) gene dataset from Lemey et al. [2009]. This dataset contains 192 HA sequences sampled from 20 localities across Eurasia between 1996 and 2005, representing a critical period in the emergence and global spread of H5N1.

Our analysis maintained consistency with the original study’s evolutionary modeling choices while adapting them for use within the structured coalescent framework. We employed an HKY+Γ substitution model, using Jeffrey’s prior [Jeffreys, 1946] on the transitiontransversion ratio and assumed an uncorrelated relaxed molecular clock model with an underlying log-normal distribution [Drummond et al., 2006]. For the discrete phylogeographic component, we implemented a normalized general substitution model [Edwards et al., 2011] with symmetric rate scaling by equilibrium frequencies. This parameterization reduces the effective number of free parameters from 380 to 190 while maintaining biological realism. We assigned gamma priors (shape = 1.0, scale = 1.0) to all migration rates, and we also constrained all 20 demes to share a single effective population size parameter. This constraint ensures compatibility between the FIT and BIT approaches while reducing parameter dimensionality [Vaughan et al., 2014]. The analysis ran for 50 million iterations with samples drawn every 10 000 iterations. Still, convergence was confirmed using Tracer v1.7 [Rambaut et al., 2018], with all key parameters achieving ESS values exceeding 200. We again constructed summary trees using the HIPSTR approach as implemented in TreeAnnotator X [Baele et al., 2024].

The computational challenges for this dataset are even more pronounced than for the dengue example. With 20 geographic locations spanning across Asia, the state space for the structured coalescent model encompasses 190 potential migration rates. Our parallel implementation reached adequate ESS values in approximately one-third of a day with two processor cores, compared to more than five days required by the BASTA package in BEAST 2.7.7 for equivalent convergence. This improvement is crucial not merely for practical feasibility but also for achieving the long runs necessary to ensure convergence and proper mixing in such a complex parameter space.

The phylogeographic reconstructions reveal markedly different patterns of uncertainty between the FIT and BIT methods (Figure 5). The FIT analysis places the root of the HA phylogeny with high confidence in Hong Kong, consistent with the 1997 outbreak that first brought H5N1 to global attention. The posterior distribution shows Hong Kong dominating with some secondary support for Guangdong and minimal probability assigned to other locations. In contrast, the BIT method exhibits substantial uncertainty regarding the root location. While Hong Kong still receives appreciable support, the posterior probability is distributed across numerous locations including Guangdong, other Chinese provinces (Fujian, Guangxi), and even more distant locations.

**Figure 5.**
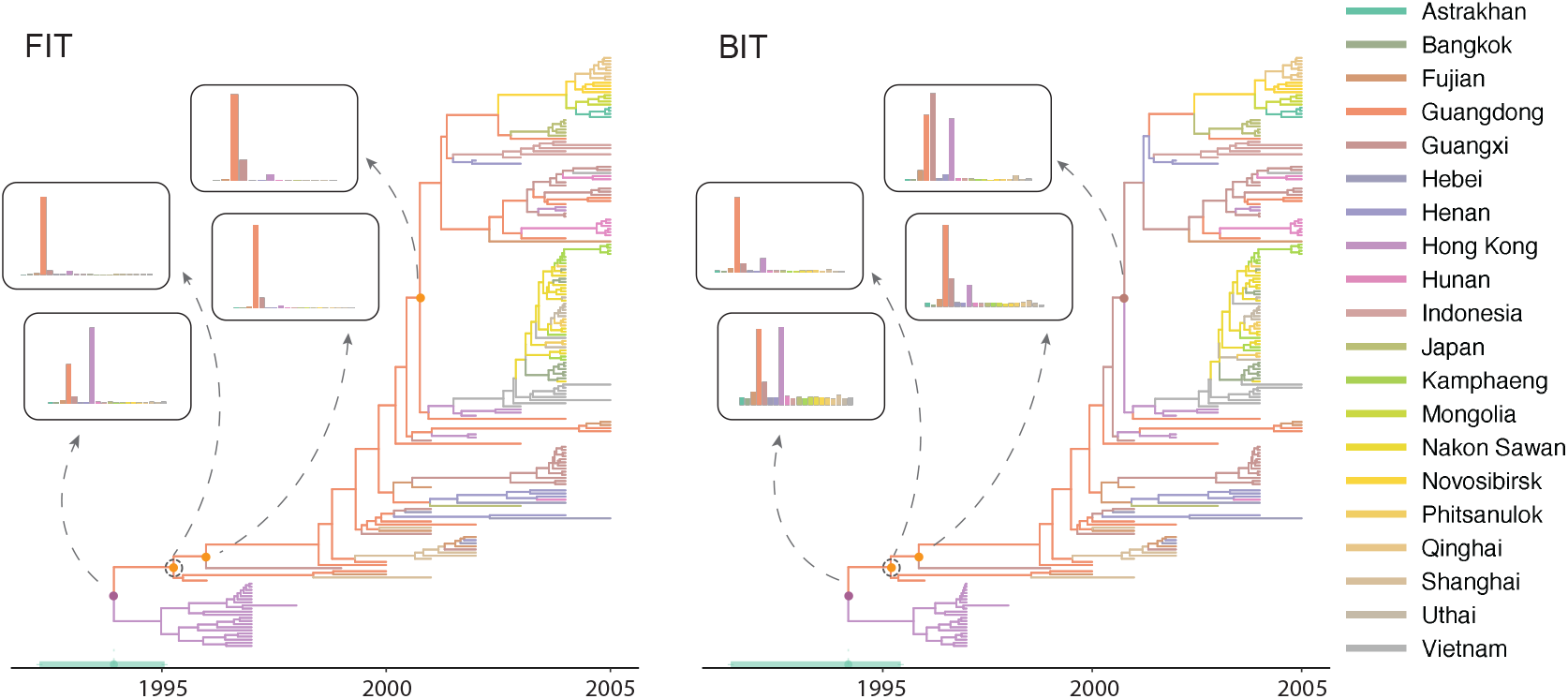
Phylogeographic reconstruction of avian influenza A H5N1 hemagglutinin (HA) across Eurasia. Highest independent posterior subtree reconstruction (HIPSTR) trees with branches colored by location state for both the FIT approach and the BIT implementation. Insets show posterior distributions of ancestral locations at key nodes, including the root and the Gs/GD lineage ancestor (marked with circles, named after A/goose/Guangdong/1/96 strain). Green bar on the time axis represents the 95% HPD credible interval for the root age.

The temporal reconstruction under both models places the root in 1994-1995, shortly before the first recognized H5N1 outbreak in Hong Kong’s poultry markets [Shortridge et al., 1998]. The branching structure reveals an initial period of limited diversity followed by rapid expansion after 2000. Both methods capture the major radiation of H5N1 lineages that occurred in the early 2000s, though they differ in some of their assignments of ancestral locations during this expansion phase. The FIT method tends to maintain strong support for southern Chinese locations (particularly Guangdong, Hong Kong and Guangxi) at internal nodes, creating a narrative of expansion from this epicenter. The BIT method, while showing the same general pattern, acknowledges considerable uncertainty about the precise geographic paths of spread.

The different uncertainty profiles between these methods likely reflect fundamental differences in how they integrate over the space of possible migration histories. The BIT approach must reconcile the observed tip locations with the coalescent process while simultaneously inferring migration along branches. This creates a more complex inference problem that naturally leads to greater uncertainty, particularly when multiple geographic scenarios could plausibly explain the observed data. The FIT approach, by contrast, conditions more directly on the reconstructed phylogeny, potentially leading to overconfidence in ancestral state assignments. De Maio et al. [2015b] previously also addressed that discrete trait models like FIT often exhibit strong overconfidence in ancestral location estimates, a key motivation for the development of the BASTA method. Acknowledging this bias underscores the importance of using approaches that properly account for migration uncertainty, even when they incur greater computational complexity.

Despite these differences in uncertainty quantification, both methods converge on key aspects of H5N1 spatial dynamics. The rapid geographic expansion after 2000 is evident in both reconstructions, with the virus spreading from southern China throughout Southeast Asia and eventually reaching Central Asia and beyond. The timing of major branching events is also consistent between methods, suggesting that while the precise geographic origins of lineages may be uncertain, the temporal dynamics of the H5N1 emergence are robustly captured.

## 5 Conclusion

We have presented a comprehensive parallel computing framework that fundamentally transforms the computational tractability of the SCA model through novel algorithmic reformulations and multi-threaded CPU implementations. Our approach achieves speedup factors ranging from 10-to 26-fold relative to the original structured coalescent implementation in BEAST 2.7.7, with optimal performance typically observed when utilizing 8-16 processor cores, contingent upon dataset characteristics. These substantial efficiency gains render previously intractable phylogeographic analyses computationally feasible, reducing requisite computation times from weeks to days for complex viral datasets.

Since optimal core utilization varies with dataset complexity, we recommend conducting short pilot runs on 8, 12, and 16 cores with fixed tree parameters to identify the most efficient configuration. BEAST X’s checkpointing functionality [Gill et al., 2020] further facilitates such optimization by enabling safe interruption and resumption of these computationally intensive analyses.

The principal innovation resides in our reformulation of the likelihood computation to expose parallelism across several computational levels: transition probability matrix calculations, partial likelihood vector propagation, and interval-wise reductions. Specifically, we implemented thread-level parallelism to distribute likelihood computations across multiple processor cores, reducing the effective time complexity from 𝒪 (*NS*^3^ + *N* ^2^*S*^2^ + *N* ^2^*S*) to approximately 𝒪 (*NS*^3^*/P* + *N* ^2^*S*^2^*/P* + *N* ^2^*S/P* ) for *P* processor cores, significantly improving scalability for high-dimensional inference problems. Furthermore, by incorporating our improvements into the BEAGLE library [Ayres et al., 2019], we enable any software leveraging this library to benefit from enhanced parallel performance with minimal additional development effort.

Our empirical applications demonstrate the transformative impact of these computtional advances on real-world phylogeographic inference. For the dengue virus serotype 1 (DENV-1) dataset comprising 287 genomic sequences distributed across 10 South American locations, our parallelized SCA implementation in BEAST X reduced requisite computation time from exceeding one week to approximately one and a half days while maintaining adequate Markov chain mixing. Notably, the SCA formulation yielded substantially more diffuse posterior distributions for ancestral location states, particularly pronounced at intermediate phylogenetic nodes, suggesting a more statistically conservative approach to uncertainty quantification relative to FIT implementations.

For the H5N1 hemagglutinin dataset encompassing 192 sequences sampled across 20 Eurasian locations, we achieved a reduction in analysis time from over five days to less than half a day. This computational efficiency enabled robust inference across the 190-dimensional migration rate parameter space, revealing the complex spatiotemporal dynamics underlying H5N1’s emergence from southern China, while appropriately quantifying substantial uncertainty regarding precise geographic transmission pathways during the early 2000s expansion phase.

Several future methodological extensions will further augment the framework’s inferential capabilities. Many-core-accelerated implementations using graphics processing units (GPUs) are under development to exploit the massive parallelization potential inherent in structured coalescent likelihood computations. The evaluation of partial likelihood vectors across nodes and the computation of interval-wise reductions naturally map onto GPU architectures, promising speedups that potentially exceed two orders of magnitude for datasets with numerous locations. For analyses involving complex migration networks or fine-scale temporal discretization, this parallelization will prove particularly transformative.

We implemented a dynamic scheduling approach to efficiently distribute computational tasks across available processor cores, enhancing load balancing in structured coalescent likelihood computations. In future work, we aim to incorporate adaptive strategies inspired by recent advances in Bayesian phylogenetic computation. Baele et al. [2017] demonstrated that combining adaptive MCMC with intelligent multi-core parallelization yielded up to 14-fold performance improvements by learning optimal parameter proposals while simultaneously distributing computational workload across processors. Building on these insights, we plan to develop adaptive mechanisms that monitor real-time computational performance and dynamically adjust resource allocation, potentially achieving more consistent parallel efficiency and reducing overall computation time in structured coalescent analyses.

Efficient gradient computation algorithms represent another key advancement, enabling Hamiltonian Monte Carlo (HMC) inference for structured coalescent approximations [Betancourt, 2017]. By analytically deriving log-likelihood gradients with respect to arbitrary dimensions of effective population sizes and migration rates through reverse-level-order traversal, the framework can efficiently explore high-dimensional parameter spaces and carry out inference under complex demographic parameterizations.

The fundamental challenge of parameter proliferation also demands innovative solutions. Similar to modelling migration rates in the FIT method as a function of a number of potential geographic dispersal covariates [Lemey et al., 2014], the SCA can also incor-porate predictor-driven migration rates that express demographic parameters as functions of observable covariates through, for example, log-linear models. This log-linear regression integration dramatically improves parameter identifiability by reducing the number of migration parameters from quadratic to linear scaling with predictors, while enabling hypothesis-driven inference about dispersal mechanisms. By incorporating external information such as geographic distances, environmental variables, or demographic factors, the framework transforms phylogeographic analysis from purely descriptive reconstruction to mechanistic understanding of pathogen spread within the structured coalescent framework.

Within the structured coalescent analysis, modeling temporal variation in effective population size is crucial for capturing realistic evolutionary dynamics. To this end, semiparametric approaches, such as piecewise-constant models, can be employed, whereby the effective population size is assumed constant within predefined time intervals but allowed to vary between them. Recent work integrates this framework into an SCA model, demonstrating its utility for joint spatial-temporal inference across multiple demes [Müller et al., 2024]. Internally, these piecewise-constant trajectories are smoothed using Gaussian Markov random field (GMRF) priors, which encourage gradual transitions between adjacent intervals and have proven effective in population dynamics reconstruction [Minin et al., 2008, Gill et al., 2012]. Importantly, our parallelized implementation supports the development of such semi-parametric demographic model extensions without compromising computational throughput.

The optimized SCA framework is fully integrated into BEAST X [Baele et al., 2025] through accompanying implementations in the BEAGLE library [Suchard and Rambaut, 2009, Ayres et al., 2019], and analyses can easily be set up using BEAUti X. This enables users to seamlessly combine it with existing features (e.g., Bayesian stochastic search variable selection for migration rates, marginal likelihood estimation) and laying the groundwork for incorporating additional extensions such as individual travel history data or Markov jump analyses in future work.

## Data Availability

All datasets, BEAST X XML configuration files, benchmark scripts, and performance measurement data used in this study are available at https://github.com/suchard-group/structured_coalescent_cpu_supplement. This repository includes the dengue virus serotype 1 (DENV-1) and avian influenza H5N1 hemagglutinin datasets, along with instructions to reproduce all computational benchmarks and phylogeographic reconstructions presented in this paper.

## Software Availability

The parallel algorithms for structured coalescent inference described in this paper have been implemented in the BEAGLE library on the basta branch at https://github.com/beagle-dev/beagle-lib/tree/basta. The corresponding BEAST X implementation is currently available in the BigFastTreeModel branch at https://github.com/beast-dev/beast-mcmc/tree/BigFastTreeModel. These branches are provided for review purposes and will be merged into the main branch following peer review. Both implementations are freely available under open-source licenses.

## 6 Competing interests

The authors declare no competing interests.

7 Acknowledgments

YS, MAS, AR and PL acknowledge support from National Institutes of Health (NIH) grants R01 AI153044 and R01 AI162611. MAS, AR and PL further acknowledge support from the Wellcome Trust (Collaborators Award 206298/Z/17/Z, ARTIC network) and the Euro-pean Research Council (grant agreement no. 725422 – ReservoirDOCS). PL acknowledges support by the Research Foundation - Flanders (‘Fonds voor Wetenschappelijk Onderzoek - Vlaanderen’, G005323N and G051322N). GB acknowledges support from the Research Foundation - Flanders (“Fonds voor Wetenschappelijk Onderzoek - Vlaanderen,” G0E1420N, G098321N), from the European Union Horizon 2023 RIA project LEAPS (grant agreement no. 101094685), and from the DURABLE EU4Health project 02/2023-01/2027 which is co-funded by the European Union (call EU4H-2021-PJ4) under Grant Agreement No. 101102733. Views and opinions expressed are however those of the author(s) only and do not necessarily reflect those of the European Union or the European Health and Digital Executive Agency. Neither the European Union nor the granting authority can be held responsible for them.

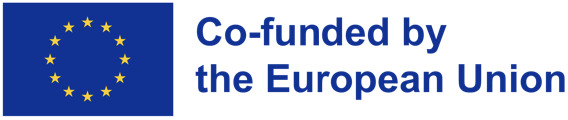

## Supplementary Materials

## Additional benchmark results

To isolate the performance gains from our algorithmic restructuring versus parallelization benefits, we performed a single-threaded comparison of structured coalescent calculations between BEAST X CPU and the BASTA package in BEAST 2.7.7. Supplementary Figure S1 demonstrates that even without any parallelization, BEAST X CPU achieves substantial speedup factors of 8.2× for EBLV (51 taxa, 3 demes), 7.0× for ZIKV (283 taxa, 22 demes), and 7.0× for PEDV (756 taxa, 26 demes). These performance improvements arise solely from our fundamental restructuring of the SCA likelihood computation, which eliminates redundant calculations through efficient caching of partial likelihoods, optimizes memory access patterns, and streamlines the peeling algorithm. The consistent 7-8× speedup across datasets of varying complexity indicates that our algorithmic optimizations provide robust baseline improvements independent of hardware parallelization. This single-threaded performance gain is particularly important as it benefits all users regardless of their computational resources, while the additional parallelization capabilities shown in the main text (Figure 1) can provide further acceleration on multi-core systems.

**Supplementary Figure S1:**
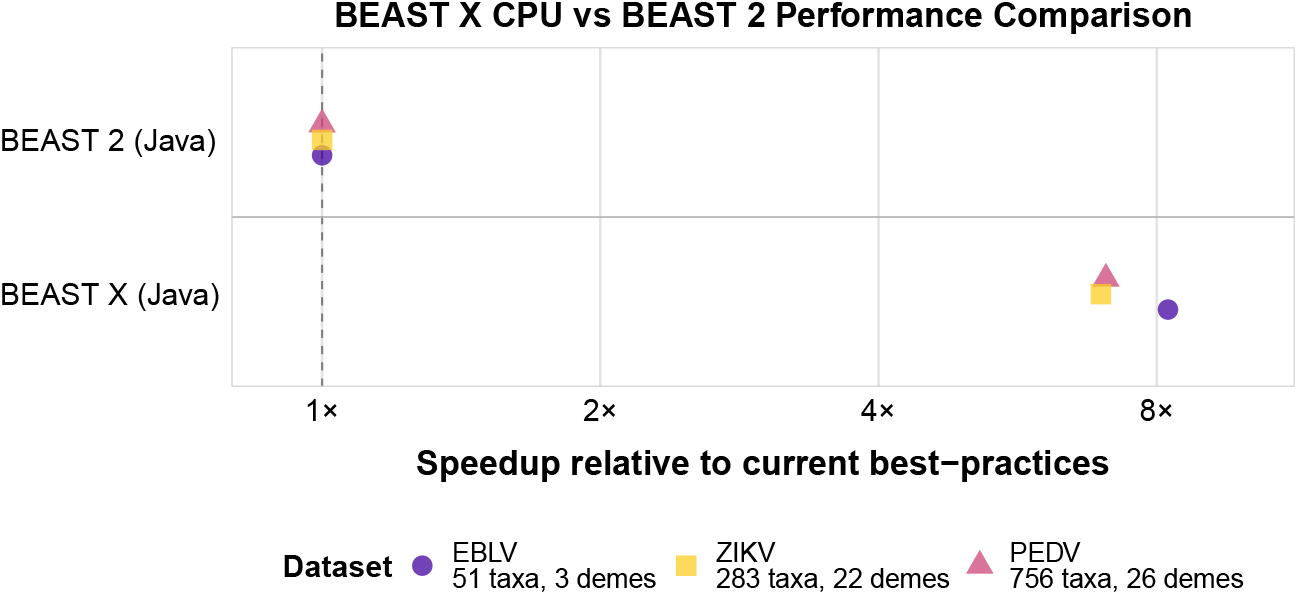
Performance comparison between single-threaded BEAST X CPU and BEAST 2.7.7 for structured coalescent approximation (SCA) analyses. The plot shows speedup factors on a log scale for three viral datasets: EBLV (51 taxa, 3 geographic states), ZIKV (283 taxa, 22 states), and PEDV (756 taxa, 26 states). BEAST X CPU achieves 7.0– 8.2× speedup over BASTA package in BEAST 2.7.7 through algorithmic restructuring alone, without any parallelization. The vertical dashed line at 1× represents baseline BEAST 2.7.7 performance.

